# MetScore and classical–basal subtype intersect to define chemotherapy response and tumor microenvironment architecture in pancreatic ductal adenocarcinoma

**DOI:** 10.64898/2026.09.14.751470

**Authors:** Raymond E. Preston, Hongyu Huang, Margaret A. Hall, Udhayvir S. Grewal, Kawther Abdilleh, Sudheer Doss, Richard Moffitt, Olatunji B. Alese, Gregory B. Lesinski, Jesse S. Handler

## Abstract

**Background:** Transcriptomic heterogeneity in pancreatic ductal adenocarcinoma (PDAC) is conventionally modeled along a single classical–basal axis, but this framework incompletely explains clinical behavior. We hypothesized that MetScore, a single-sample measure of metastatic colonization potential, defines a complementary axis.

**Objective:** To determine how MetScore and classical–basal identity jointly shape PDAC biology and clinical outcomes.

**Design:** We integrated RNA sequencing, DNA sequencing, clinical, and digital pathology data from 512 tumor biopsies from 510 patients in the Pancreatic Cancer Action Network’s Know Your Tumor real-world cohort. We assessed associations with biopsy site, survival, chemotherapy response, and tumor microenvironment composition; validated metastatic discrimination in an independent cohort; and examined cancer cell–intrinsic drug sensitivity *in vitro*.

**Results:** MetScore distinguished metastatic from primary biopsies, a finding validated externally. High MetScore and basal identity were independently associated with inferior survival. High MetScore and classical identity were each associated with preferential response to 5-fluorouracil (5-FU)–based rather than gemcitabine-based therapy, with the strongest enrichment for 5-FU responses among classical/MetScore-high patients. *In vitro* drug-sensitivity experiments did not recapitulate the clinical treatment-response pattern, motivating investigation of cancer cell–extrinsic factors. Transcriptomic deconvolution and histopathologic machine learning-based cell annotation demonstrated that high-MetScore and basal states converged on macrophage/monocyte enrichment but diverged in their associations with cancer-associated fibroblasts.

**Conclusion:** Integrating MetScore with classical–basal identity establishes a two-axis framework that better explains metastatic behavior, prognosis, treatment response, and microenvironmental composition than the conventional one-axis model and provides a clinically testable strategy for transcriptome-guided patient stratification.

## INTRODUCTION

Transcriptomic state, the coordinated pattern of messenger RNA expression that reflects a cell’s functional phenotype, is emerging as a clinically important dimension of cancer biology. Distinct gene expression programs are associated with key properties of tumor progression, including therapy resistance^1-3^, immune evasion^4^, and metastatic competence^5-7^. Because these programs capture biological phenotypes that may not be evident from DNA sequence alterations alone, transcriptomic biomarkers have the potential to complement DNA profiling and improve patient stratification.

In pancreatic ductal adenocarcinoma (PDAC), transcriptomic studies have consistently identified two dominant tumor cell-intrinsic programs: classical and basal^8-10^. These molecular subtypes are strongly prognostic, with basal PDAC consistently associated with shorter survival. This classical–basal axis has consequently emerged as the principal framework for describing malignant-cell differentiation in PDAC and forms the basis of clinically applicable classifiers such as PurIST^11^.

Classical–basal differentiation has also been associated with chemotherapy response and tumor microenvironment (TME) composition. Retrospective clinical studies have suggested that classical tumors respond more favorably to FOLFIRINOX and may derive greater benefit from FOLFIRINOX than from gemcitabine plus nab-paclitaxel, whereas basal tumors generally respond poorly to chemotherapy^11 12^. However, the randomized PASS-01 trial did not identify significant progression-free survival differences between these regimens within either molecular subtype^13^, indicating that classical–basal subtype alone may be insufficient to guide chemotherapy selection. Basal malignant programs have been linked to activated fibroinflammatory microenvironments enriched for myofibroblastic cancer-associated fibroblasts (CAFs), SPP1-positive tumor-associated macrophages (TAMs), and regulatory T cells^14^. These observations establish the clinical and biological importance of classical–basal differentiation while suggesting that it does not capture the full spectrum of transcriptomic variation relevant to PDAC behavior.

We recently identified a distinct metastatic-potential cell-state axis in PDAC through integrated analyses of bulk and single-cell genomic datasets^5^. This axis, quantified by MetScore, was associated with metastatic colonization and adverse clinical outcomes and was orthogonal to classical–basal differentiation. Thus, metastatic potential and classical–basal subtype represent independent dimensions of PDAC transcriptomic state. However, how these axes intersect to modulate chemotherapy response and tumor microenvironmental architecture remained unknown.

Here, we used large real-world clinicogenomic cohorts, transcriptome-based microenvironmental inference, digital pathology, and experimental models to define the clinical and biological associations of these two transcriptomic axes. We validated MetScore across independent patient cohorts and commercial sequencing platforms and found that it was associated with systemic, but not transcoelomic, dissemination and provided prognostic information independent of classical–basal subtype. MetScore also modified the association between chemotherapy backbone and objective response, with classical/MetScore-high tumors exhibiting the greatest response enrichment with 5-FU-based therapy. Finally, the two axes showed opposing associations with fibroblast abundance but concordant associations with macrophage/monocyte enrichment. Together, these findings establish a two-axis framework for understanding clinically relevant transcriptomic heterogeneity in PDAC, provide a strategy for transcriptome-based patient stratification, and identify aggressive PDAC cell states for therapeutic targeting.

## MATERIALS AND METHODS

Detailed description is included in the supplementary materials.

## RESULTS

### The MetScore-defining cell-state axis is associated with systemic but not transcoelomic dissemination in PDAC

In our previous study^5^, MetScore, a scoring metric capturing a metastatic-potential cell-state axis, distinguished metastatic PDAC tumors and cells from their primary-tumor counterparts in the Moffitt^8^ and ICGC^15 16^ bulk transcriptomic cohorts and in a single-cell RNA-sequencing atlas^17^. However, whether this association generalizes to real-world patient populations and commercial next-generation sequencing platforms remained unknown. We therefore analyzed the Pancreatic Cancer Action Network Know Your Tumor (KYT) cohort, which includes clinical data and matched tumor transcriptomic profiles generated using the Tempus platform from 510 patients with PDAC treated in community settings (**figure 1 and table 1**). Although genomic findings from KYT have previously been reported^18 19^, the transcriptomic features of this cohort have not been comprehensively characterized.

**Table 1.** Patient characteristics.

| Patient characteristics |  |
| --- | --- |
| Characteristic | Evaluable cohort<br>(n = 510)<br>N (%) |
| Sex |  |
| Male | 260 (51.0%) |
| Female | 250 (49.0%) |
| Age at diagnosis |  |
| < 50 years | 25 (4.9%) |
| 50-54 years | 25 (4.9%) |
| 55-59 | 55 (10.8%) |
| 60-64 | 58 (11.4%) |
| 65-69 | 64 (12.5%) |
| 70-74 | 38 (7.5%) |
| 75-79 | 26 (5.1%) |
| ≥ 80 | 8 (1.6%) |
| Missing | 211 (41.4%) |
| Race |  |
| White | 134 (26.3%) |
| Black or African American | 10 (2.0%) |
| Asian | 9 (1.8%) |
| Other Race | 5 (1.0%) |
| Missing | 352 (69.0%) |
| Stage at diagnosis |  |
| M 0 | 234 (45.9%) |
| M 1 | 276 (54.1%) |

**Figure 1.**
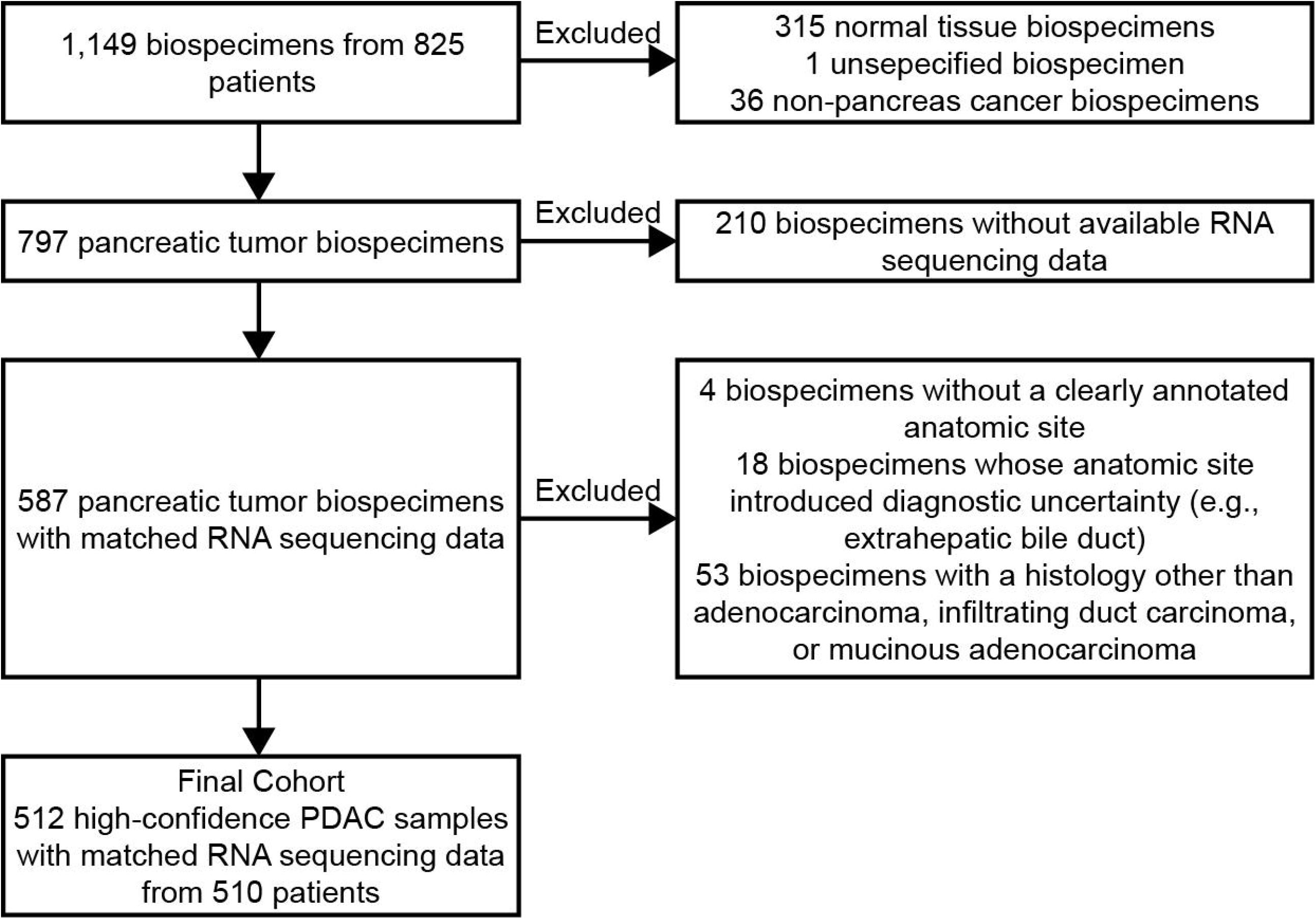
The Know Your Tumor cohort diagram.

Within the KYT analytic cohort, metastatic-site biopsies (n = 194) had higher MetScores than primary-tumor biopsies (n = 318; Wilcoxon rank-sum test *p* < 0.001; **figure 2A**). To assess reproducibility in an independent real-world cohort profiled using a different commercial platform, we analyzed patients with PDAC treated at the Winship Cancer Institute of Emory University whose tumors underwent molecular profiling using the Caris Life Sciences platform between 2024 and 2025. MetScores were again higher in metastatic-site biopsies (n = 58) than in primary-tumor biopsies (n = 84; Wilcoxon rank-sum test *p* = 0.017; **figure 2B**). Thus, the association between MetScore and metastatic biopsy site was reproducible across independent clinical cohorts and commercial sequencing platforms.

**Figure 2.**
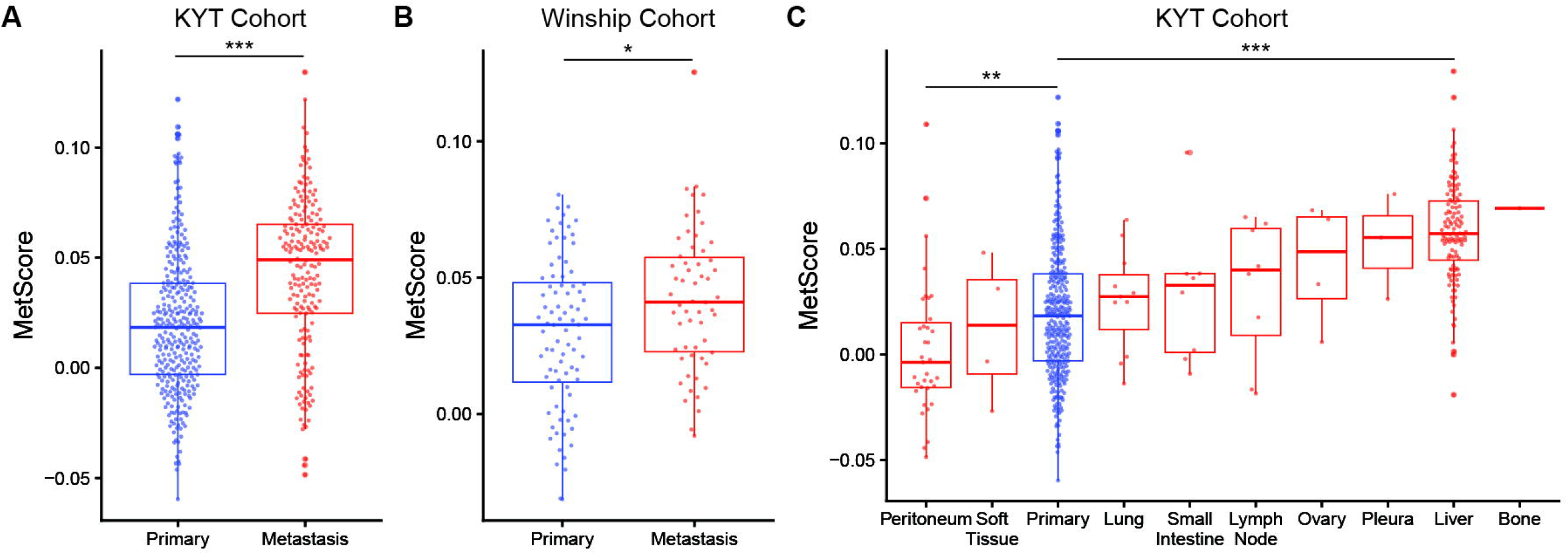
The MetScore-defining cell-state axis is associated with systemic but not transcoelomic dissemination in PDAC. **(A)** MetScore distributions in primary-tumor and metastatic-site biopsies from the Know Your Tumor (KYT) cohort. **(B)** MetScore distributions in primary-tumor and metastatic-site biopsies from the independent Winship cohort. **(C)** MetScore distributions in the KYT cohort after stratification of metastatic biopsies by anatomic site. Primary and metastatic-site biopsies in **A** and **B** were compared using two-sided Wilcoxon rank-sum tests. In **C**, each metastatic site was compared with primary tumors using two-sided Wilcoxon rank-sum tests, with Benjamini–Hochberg correction for multiple comparisons; only liver and peritoneal metastases remained significantly different from primary tumors after correction. Boxes indicate the median and interquartile range; whiskers extend to the most extreme values within 1.5 times the interquartile range of the hinges. Each point represents one tumor biopsy. \**P* < 0.05, \*\**P* < 0.01, and \*\*\**P* < 0.001. Significance symbols in C denote Benjamini–Hochberg-adjusted *P* values.

We next examined whether the association between MetScore and metastasis varied across sites characterized by different routes of dissemination. Among non-peritoneal metastatic sites represented by at least five biopsies, each had a higher mean MetScore than primary-tumor biopsies (**figure 2C**). Liver metastases were the most frequently sampled (n = 120), had the highest mean MetScore among these sites, and had significantly higher MetScores than primary-tumor biopsies (Wilcoxon rank-sum test Benjamini-Hochberg corrected *p* < 0.001). In contrast, peritoneal metastases (n = 35) had lower MetScores than primary-tumor biopsies (*p* = 0.004). Collectively, these findings indicate that high MetScore is associated with dissemination to non-peritoneal sites but not with peritoneal metastasis, consistent with biological differences between vascular or lymphatic dissemination and transcoelomic spread.

### Higher MetScore and basal subtype are independently associated with shorter overall survival in PDAC

In our previous study^5^, MetScore and classical–basal subtype provided nonredundant prognostic information. However, these findings were derived from relatively small academic cohorts, and their generalizability to the broader PDAC population remained uncertain. We therefore applied the PurIST classifier^11^ to assign each tumor in the KYT cohort a classical or basal subtype and evaluated the associations of MetScore and subtype with overall survival among patients with both transcriptomic and clinical outcomes data (n = 299).

In a Cox proportional hazards model containing both molecular features, each one standard deviation increase in MetScore was associated with a 55% higher hazard of death (HR 1.55, 95% CI 1.32–1.82; *p* < 0.001). Independently, basal subtype was associated with shorter overall survival compared with classical subtype (HR 1.81, 95% CI 1.27–2.58; *p* = 0.001). To visualize the joint relationship of these molecular features with survival, we dichotomized MetScore at the cohort median and divided patients into four groups: classical/MetScore-low, classical/MetScore-high, basal-like/MetScore-low, and basal-like/MetScore-high. Kaplan–Meier estimates showed that patients with classical/MetScore-low tumors had the longest overall survival, whereas those with basal-like/MetScore-high tumors had the shortest; the classical/MetScore-high and basal-like/MetScore-low groups had overlapping intermediate outcomes (overall log-rank *p* < 0.001; **figure 3A**).

**Figure 3.**
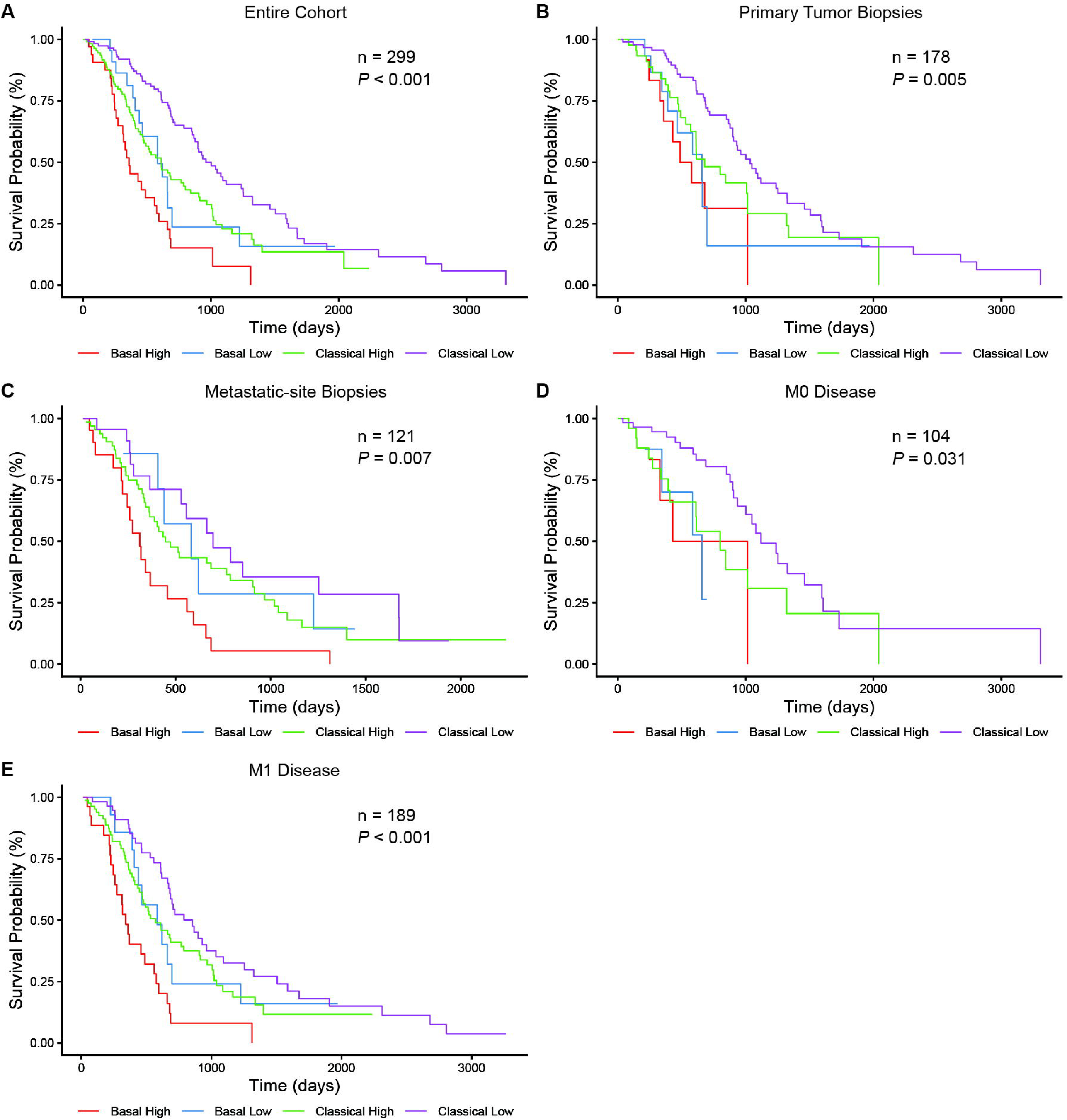
Higher MetScore and basal subtype are independently associated with shorter overall survival in PDAC. Kaplan–Meier estimates of overall survival among patients in the Know Your Tumor (KYT) cohort, divided into four molecular groups according to classical–basal subtype determined using PurIST and MetScore dichotomized at the cohort median: classical/MetScore-low, classical/MetScore-high, basal-like/MetScore-low, and basal-like/MetScore-high. (**A**) Entire survival-evaluable cohort. (**B, C**) Patients stratified by biopsy site: primary-tumor biopsies (**B**) and metastatic-site biopsies (**C**). (**D, E**) Patients stratified by metastatic status at diagnosis: M0 disease (**D**) and M1 disease (**E**). Differences among the four molecular groups were evaluated using two-sided log-rank tests. Sample sizes and overall log-rank *P* values are shown in each panel. Tick marks indicate censored observations.

To assess the robustness of these associations across clinically relevant subgroups, we repeated the Cox models after stratification by biopsy site and stage at diagnosis. Among patients with primary tumor biopsies (n = 178), each one standard deviation increase in MetScore (HR 1.37, 95% CI 1.08–1.73; *p* = 0.010) and basal subtype (vs classical; HR 1.88, 95% CI 1.11–3.20; *p* = 0.020) were associated with shorter overall survival (**figure 3B**). Similar associations were observed among patients with metastatic-site biopsies (n = 121) for both MetScore (HR 1.50, 95% CI 1.18–1.91; *p* < 0.001) and basal subtype (HR 1.64, 95% CI 1.01– 2.65; *p* = 0.046; **figure 3C**).

When stratified by stage at diagnosis, higher MetScore remained associated with shorter overall survival in both patients with M0 disease (n = 104; HR per 1-SD increase 1.56, 95% CI 1.09– 2.22; *p* = 0.015) and those with M1 disease (n = 189; HR 1.45, 95% CI 1.20–1.74; *p* < 0.001; **figures 3D,E**). Basal subtype had directionally similar associations in the M0 (HR 2.14, 95% CI 0.96–4.77; *p* = 0.063) and M1 subgroups (HR 1.76, 95% CI 1.18–2.63; *p* = 0.005; **figures 3D,E**), although the estimate in the smaller M0 subgroup was less precise. Formal interaction testing provided no evidence that the associations of MetScore or subtype with overall survival differed by biopsy site (MetScore, *p* interaction = 0.29; subtype, *p* interaction = 0.81) or metastatic status at diagnosis (MetScore, *p* interaction = 0.47; subtype, *p* interaction = 0.96).

Collectively, these findings support the independent prognostic relevance of MetScore and basal subtype in a large, community-based PDAC cohort.

### Classical/MetScore-high PDAC is associated with a higher response rate to 5-FU-than gemcitabine-based therapy

Prior transcriptomic studies associated classical PDAC with more favorable outcomes following multiagent chemotherapy than basal PDAC, raising the possibility that molecular subtyping could inform chemotherapy selection. However, the randomized PASS-01 trial did not establish a clear treatment-selection strategy based on classical–basal subtype alone. We therefore asked whether integrating MetScore with molecular subtype could identify PDAC states associated with differential responses to chemotherapy in the KYT cohort.

We identified 112 patients with transcriptomic profiles and linked treatment and response data. Given the heterogeneity of treatments administered in this real-world cohort, regimens were classified according to whether they contained a 5-FU or gemcitabine backbone using prespecified criteria (see **Materials and Methods**). Regimens containing both backbones or those that could not be unambiguously classified were excluded. To limit confounding from prior therapy, the analysis was restricted to patients receiving first-line treatment, resulting in a final response-evaluable cohort of 97 patients. Objective response rate (ORR) was defined as the proportion of patients whose best documented response was a complete or partial response.

Among patients with MetScore-high tumors, ORR was higher with 5-FU-than with gemcitabine-based therapy (20/34 [59%] vs 5/22 [23%]; OR 4.71, 95% CI 1.28–20.4; Fisher’s exact *p* = 0.013). No treatment-associated difference was detected among patients with MetScore-low tumors (OR 0.61, 95% CI 0.13–2.65; *p* = 0.520; **figure 4A**). Consistent with these subgroup findings, there was evidence of a treatment-by-MetScore interaction in logistic regression (interaction *p* = 0.021), indicating that the association between chemotherapy backbone and objective response differed by MetScore group.

**Figure 4.**
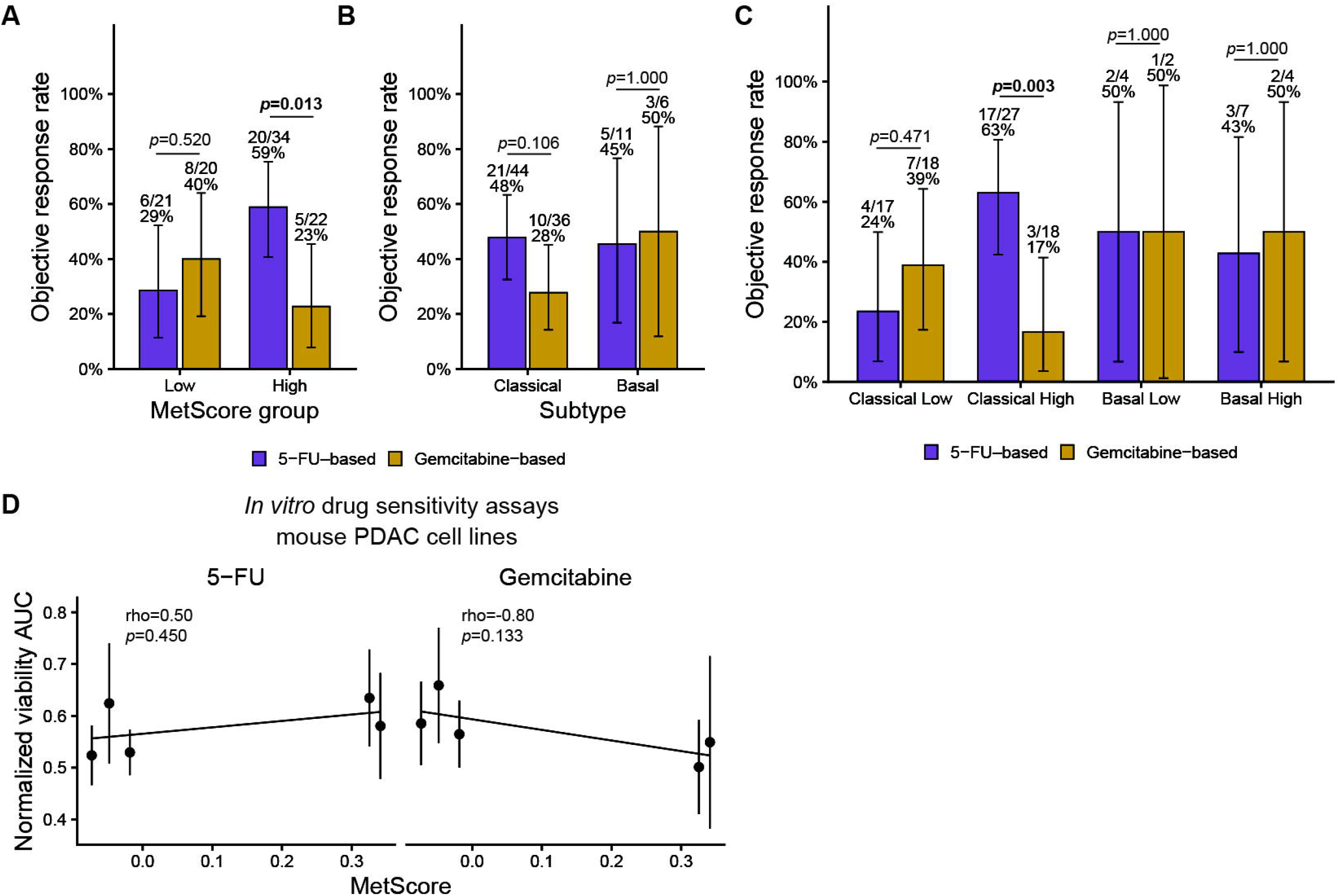
Classical/MetScore-high PDAC is associated with a higher objective response rate to 5-FU-than gemcitabine-based therapy. (**A-C**) Objective response rates (ORRs) according to chemotherapy backbone and molecular state in the Know Your Tumor (KYT) cohort. **(A)** ORR with 5-FU- and gemcitabine-based therapy among patients with MetScore-low or MetScore-high tumors, defined by dichotomization at the cohort median. **(B)** ORR according to chemotherapy backbone among patients with classical or basal tumors, as determined using PurIST. **(C)** ORR according to chemotherapy backbone after joint stratification by MetScore and classical–basal subtype. The analysis included patients receiving first-line therapy whose regimen could be unambiguously classified as 5-FU- or gemcitabine-based and for whom pretreatment RNA-sequencing and evaluable response data were available (n = 97). Bars represent ORRs, and error bars indicate 95% binomial confidence intervals. The number of responders, total number of patients, and ORR are displayed for each treatment group. Within each molecular group, ORRs with 5-FU- and gemcitabine-based therapy were compared using two-sided Fisher’s exact tests. Reported *P* values are nominal and were not adjusted for multiple comparisons. **(D)** Relationship between MetScore and normalized viability area under the curve (AUC) for 5-fluorouracil (5-FU; left) and gemcitabine (right) in five murine PDAC cell lines. Points represent individual cell lines (mean ± SEM from three independent experiments). Lines show linear regression fits. Spearman’s correlation coefficients (ρ) and exact two-sided *P* values were calculated across the five cell-line means.

Among patients with classical tumors, ORR was numerically higher with 5-FU-than with gemcitabine-based therapy, consistent with prior reports^11 12^, although the difference was not statistically significant (21/44 [48%] vs 10/36 [28%]; OR 2.35, 95% CI 0.85–6.84; Fisher’s exact *p* = 0.106). No treatment-associated difference was detected among patients with basal tumors (OR 0.84, 95% CI 0.07–9.45; *p* = 1.000; **figure 4B**). The treatment-by-subtype interaction did not reach significance (interaction *p* = 0.352).

Because the point estimates for both high MetScore and classical subtype favored 5-FU-based therapy, we next examined whether their intersection defined a 5-FU response-enriched molecular state. Patients with classical/MetScore-high tumors had a higher ORR with 5-FU-than with gemcitabine-based therapy (17/27 [63%] vs 3/18 [17%]; OR 8.06, 95% CI 1.70–54.3; Fisher’s exact *p* = 0.003; **figure 4C**). No statistically significant treatment-associated differences were detected in the other joint molecular groups. Thus, classical/MetScore-high tumors exhibited the largest observed treatment-associated difference in ORR, although allowing treatment effects to vary across the four combined MetScore/subtype groups did not significantly improve model fit relative to a model in which treatment effects varied by MetScore alone (*p* = 0.237). Collectively, these findings identify MetScore as a correlate of differential response to 5-FU- and gemcitabine-based therapy and nominate classical/MetScore-high PDAC as a particularly 5FU response-enriched molecular state.

We next asked whether the preferential response to 5-FU observed among classical/MetScore-high tumors reflected differences in cancer cell–intrinsic sensitivity to 5-FU or gemcitabine. We generated dose–response curves in five previously established murine classical PDAC cell lines with a range of MetScores. Correlations between MetScore and normalized viability AUC did not reach statistical significance following either 5-FU (Spearman ρ = 0.50, *p* = 0.450) or gemcitabine treatment (ρ = −0.80, *p* = 0.133; **figure 4D**). Thus, the *in vitro* results did not recapitulate the clinical treatment-response pattern, prompting us to investigate potential contributions from the TME.

### MetScore-high and basal states have opposing associations with fibroblast abundance but concordant associations with monocytic-lineage cells

To explore how the metastatic-potential and classical–basal axes jointly relate to TME composition, we first applied ESTIMATE^20^ to the KYT transcriptomic dataset to derive expression-based tumor purity, immune and stromal scores. We combined these measures into an immune–stromal balance score (see **Materials and Methods**), with higher values indicating a stronger immune relative to nonimmune stromal expression signal. MetScore was positively correlated with the immune–stromal balance score in both primary tumors (Spearman’s ρ = 0.52, *p* < 0.001) and metastatic-site biopsies (ρ = 0.53, *p* < 0.001; **figure 5A**), extending our previous single-cell observations to a larger real-world cohort and to metastatic tumors. In contrast, basal-state probability was negatively correlated with the immune–stromal balance score in primary tumors (ρ = -0.15, *p* = 0.006) and showed a weaker, nonsignificant association in metastatic-site biopsies (ρ = -0.04, *p* = 0.573; **figure 5B**). Thus, the two transcriptomic axes showed distinct associations with the relative immune and stromal signals captured by ESTIMATE.

**Figure 5.**
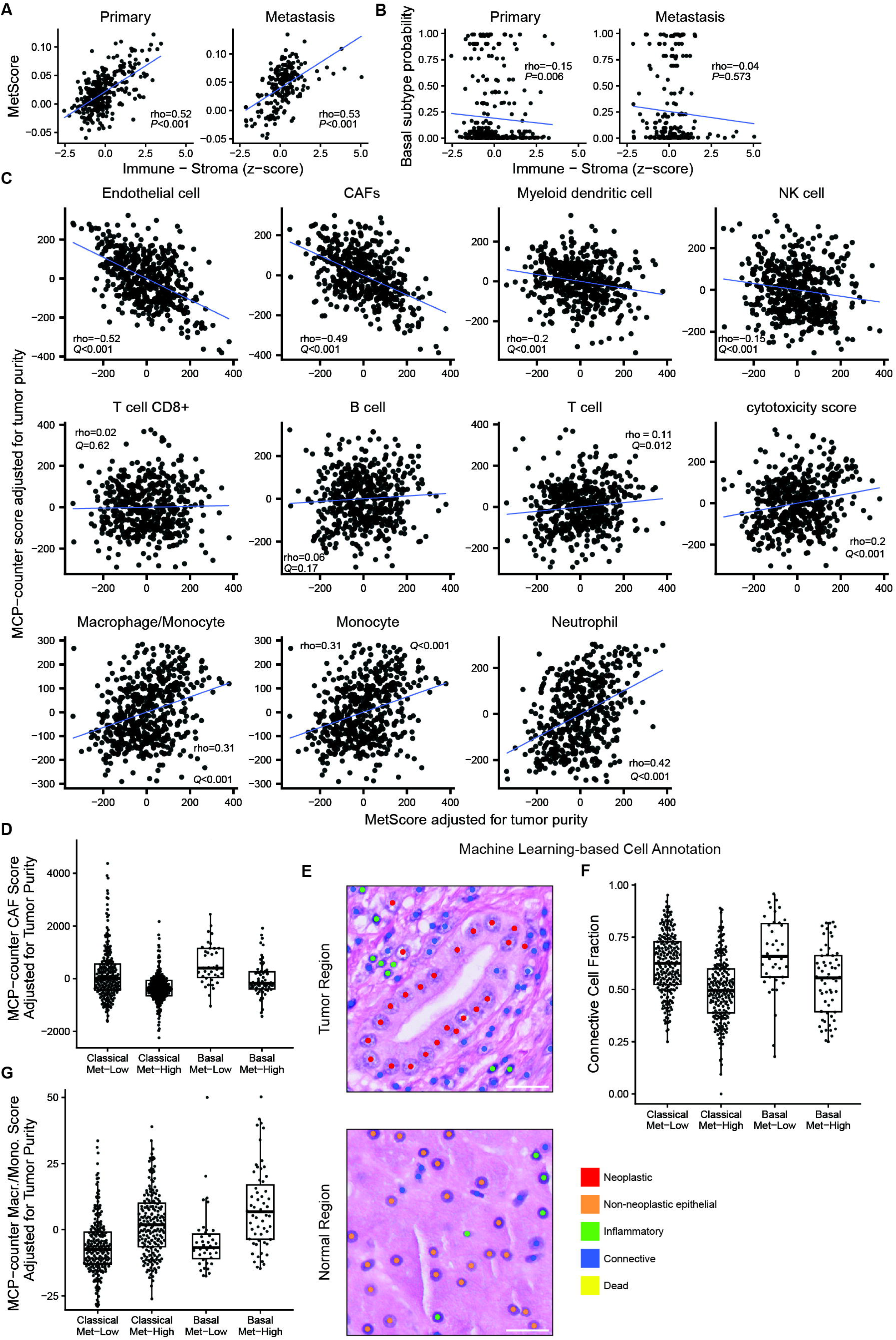
MetScore-high and basal states have opposing associations with fibroblast abundance but concordant associations with monocytic-lineage cells. Analyses in **A–D** and **G** included all tumors in the Know Your Tumor (KYT) cohort with available transcriptomic data (n = 512). Digital-pathology analysis in **E** and **F** was restricted to tumors with matched hematoxylin and eosin (H&E)-stained images (n = 502). (**A, B**) Associations of MetScore (**A**) and basal-state probability, quantified using the PurIST score (**B**), with the immune–stroma balance score. The immune–stroma balance score was calculated as the Z-standardized ESTIMATE immune score minus the Z-standardized ESTIMATE stromal score, with higher values indicating greater relative immune representation. Each point represents one tumor biopsy; primary-tumor and metastatic-site biopsies are shown separately. Least-squares regression lines are displayed for visualization. Spearman’s ρ and corresponding *P* values were calculated separately by biopsy site and are shown. **(C)** Associations between MetScore and MCP-counter cell-population scores. MetScore and MCP-counter scores were residualized for ESTIMATE-derived tumor purity before analysis. Each point represents one tumor biopsy. Least-squares regression lines are displayed for visualization, and Spearman’s ρ and corresponding *P* values are shown. (**D, F, G**) Tumors were divided into four molecular groups according to classical–basal subtype determined using PurIST and MetScore dichotomized at the cohort median: classical/MetScore-low, classical/MetScore-high, basal-like/MetScore-low, and basal-like/MetScore-high. **(D)** Tumor purity-adjusted MCP-counter fibroblast scores across the four molecular groups. Samples with scores below the 1^st^ and above the 99^th^ percentile were omitted for visualization only. **(E)** Representative images and corresponding CellViT cell-classification calls from tumor-involved and normal regions of a KYT H&E-stained tissue section. **(F)** Fraction of classified cells assigned to the CellViT connective-tissue category across the four molecular groups. **(G)** Tumor purity-adjusted MCP-counter macrophage/monoctye scores across the four molecular groups. In **D, F**, and **G**, boxes indicate the median and interquartile range; whiskers extend to the most extreme values within 1.5 times the interquartile range of the hinges. Each point represents one tumor biopsy.

To examine specific stromal populations, we next applied MCP-counter^21^ to the KYT dataset to derive transcriptome-based abundance scores. Associations were evaluated after adjustment for ESTIMATE-derived tumor purity (see **Materials and Methods**). MetScore was inversely associated with the fibroblast abundance score across the entire cohort (Spearman’s ρ = -0.49, Benjamini-Hochberg adjusted *p* < 0.001; **figure 5C**) and within both primary tumors (ρ = -0.36, *padj* < 0.001; **figure S1A**) and metastatic-site biopsies (ρ = -0.47, *padj* < 0.001; **figure S1B**). Conversely, basal-state probability was positively associated with fibroblast abundance across the entire cohort (ρ = 0.23, *padj* < 0.001; **figure S2**), primary tumors (ρ = 0.34, *padj* < 0.001; **figure S3A**), and metastatic-site biopsies (ρ = 0.18, *padj* = 0.063; **figure S3B**), although these associations were weaker than those observed for MetScore.

For visualization of their joint relationship with fibroblast abundance, tumors were divided into four molecular groups using classical–basal subtype and the median MetScore. Basal-like/MetScore-low tumors had the highest fibroblast abundance, whereas classical/MetScore-high tumors had the lowest (**figure 5D**). In a multivariable model containing continuous MetScore and basal-state probability, MetScore remained inversely associated with fibroblast abundance (standardized β = −0.37, 95% CI −0.45 to −0.29; *p* < 0.001), whereas basal-state probability remained positively associated with fibroblast abundance (standardized β = 0.25, 95% CI 0.16–0.33; *p* < 0.001). No evidence of an interaction between MetScore and basal-state probability was detected (interaction *p* = 0.17), indicating their effects were additive. These results support opposing associations of the two transcriptomic axes with fibroblast abundance.

We sought to validate this relationship at the single cell level by applying machine learning-based cell segmentation and classification using CellVit^22^ to KYT samples with available high-resolution H&E images (n = 502). CellViT classified cells as neoplastic, non-neoplastic epithelial, inflammatory, connective/soft-tissue, or dead (**figure 5E**). Classical/MetScore-high tumors had the lowest proportion of connective/soft-tissue cells among the four molecular groups and basal/MetScore-low tumors the highest (**figure 5F**), concordant with their transcriptome-derived fibroblast scores. In a multivariable model, MetScore was inversely associated with connective cell fraction (standardized β = −0.31, 95% CI −0.39 to −0.23; *p* < 0.001), whereas basal-state probability was positively associated with connective cell fraction (standardized β = 0.17, 95% CI 0.08–0.26; *p* < 0.001). Because the CellViT connective/soft-tissue category is not specific to fibroblasts and can include other mesenchymal cell populations, these data provide orthogonal support for biased nonimmune stromal cellularity rather than fibroblast-specific validation.

The two transcriptomic axes showed a different relationship with the MCP-counter macrophage/monocyte score. MetScore was positively correlated with macrophage/monocyte score across the entire cohort (Spearman ρ = 0.31, Benjamini-Hochberg adjusted *p* < 0.001; **figure 5C**) and within primary tumors (ρ = 0.15, *padj* = 0.009; **figure S1A**) and metastatic-site biopsies (ρ = 0.28, *padj* < 0.001; **figure S1B**). Basal-state probability was also positively correlated with macrophage/monocyte score in the overall cohort (ρ = 0.17, *padj* < 0.001; **figure S2**), primary tumors (ρ = 0.14, *padj* = 0.049; **figure S3A**), and metastatic-site biopsies (ρ = 0.089, *padj* = 0.27; **figure S3B**), although the associations were weaker. Basal-like/MetScore-high tumors had the highest monocytic-lineage abundance, whereas classical/MetScore-low tumors had the lowest (**figure 5G**).

In a multivariable model containing both continuous molecular scores, MetScore remained positively associated with macrophage/monocyte score (standardized β = 0.31, 95% CI 0.22 to 0.39; *p* < 0.001). The association with basal-state probability remained positive but was not statistically significant after adjustment for MetScore (β = 0.049, 95% CI -0.039 to 0.14; *p* = 0.27). The MetScore-by-basal-state interaction was not significant (interaction *p* = 0.39). Thus, MetScore showed an independent positive association with macrophage/monocyte score, whereas basal-state probability showed a directionally concordant but weaker association.

Collectively, these analyses demonstrate that the metastatic-potential and classical–basal axes are associated with distinct features of PDAC TME architecture across primary and metastatic-site biopsies

### MetScore is higher in *KRAS*-mutant and *MTAP*-deleted PDAC

To examine the relationship between MetScore, classical–basal subtype, and clinically relevant genomic biomarkers, we first compared tumors in the KYT cohort according to *KRAS* mutation status. *KRAS* mutations were detected in 88.6% of evaluable samples (442/499; see **Materials and Methods**), with the distribution of specific alleles following the expected pattern^23^ (**figure 6A**). *KRAS*-mutant tumors had significantly higher MetScores than *KRAS*-wild-type tumors (median [IQR], 0.028 [0.050] vs -0.00051 [0.051]; Wilcoxon rank-sum *p* < 0.001; **figure 6B**). When tumors were dichotomized at the median MetScore, *KRAS*^*G12D*^ was more frequent among MetScore-high than MetScore-low tumors (101/249 [41%] vs 73/250 [29%]; OR 1.7, 95% CI 1.1–2.4; Benjamini–Hochberg-adjusted Fisher’s exact *p* = 0.034; **figure 6C**). Conversely, *KRAS*-wild-type tumors were less frequent in the MetScore-high group (16/249 [6%] vs 41/250 [16%]; OR 0.35, 95% CI 0.18–0.66; adjusted *p* = 0.005). The frequencies of the other detected *KRAS* alleles did not differ significantly between MetScore groups after correction for multiple testing. Thus, MetScore-high PDAC was enriched for *KRAS* mutations, with G12D accounting for the largest component of this association. The fraction of *KRAS* WT and mutant samples classified as basal subtype was similar (10/57 (17.5%) of WT samples; 86/442 (19.5%) or mutant samples; Fisher’s exact *p* = 0.86; **figure 6D**).

**Figure 6.**
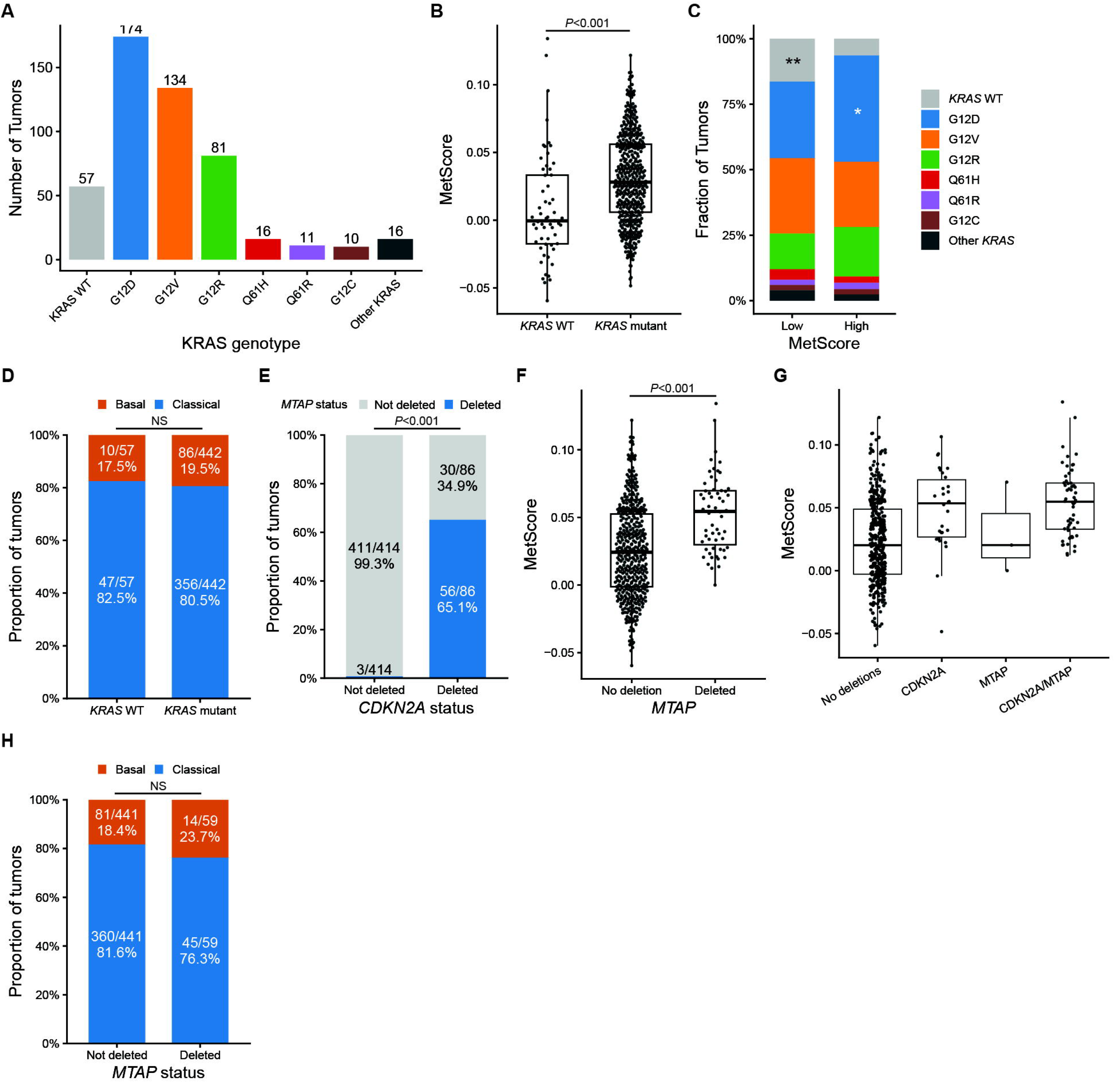
MetScore is higher in *KRAS*-mutant and *MTAP*-deleted PDAC. (**A–D**) KYT tumors with available RNA sequencing and SNV/indel calls (n = 499). **(A)** Number of tumors harboring each *KRAS* allele. Tumors without a detected *KRAS* mutation are labeled wild-type (WT); mutations not displayed separately are grouped as “Other *KRAS*.” Three tumors with both hotspot and non-hotspot variants were categorized by the hotspot mutation. **(B)** MetScore according to *KRAS* mutation status (Wilcoxon rank-sum test). **(C)** *KRAS* allele distributions in MetScore-low and MetScore-high tumors, dichotomized at the cohort median. Each allele was compared with all other categories combined using Fisher’s exact test, with Benjamini–Hochberg correction across comparisons. **(D)** PurIST classical and basal subtype frequencies according to *KRAS* mutation status (Fisher’s exact test). (**E–H**) KYT tumors with available RNA sequencing and *MTAP/CDKN2A* copy-number calls (n = 500). **(E)** Frequency of *MTAP* deletion according to *CDKN2A* deletion status (Fisher’s exact test). **(F)** MetScore according to *MTAP* deletion status (Wilcoxon rank-sum test). **(G)** MetScore across tumors with neither deletion, *CDKN2A* deletion only, *MTAP* deletion only, or *MTAP/CDKN2A* co-deletion. **(H)** PurIST classical and basal subtype frequencies according to *MTAP* deletion status (Fisher’s exact test). For (**B, F**, and **G**), boxes show medians and interquartile ranges, whiskers extend to 1.5 times the interquartile range, and points represent individual tumor biopsies. All tests were two-sided. \**P* < 0.05; \*\**P* < 0.01; NS, not significant.

Beyond *KRAS* mutation, another emerging genomic biomarker in PDAC is *MTAP* deletion, which predicts sensitivity to PRMT5 inhibition in a synthetic lethality relationship^24^. *MTAP* deletion was detected in 11.8% of evaluable samples (59/500; see **Materials and Methods**), comparable to prior population estimates^25^. *MTAP* deletion is thought to be a bystander event, frequently occurring in the context of deletion of *CDKN2A*, a tumor suppressor and nearby gene. Concordant with this model, *MTAP* deletions in the KYT cohort were highly enriched in *CDKN2A* deleted samples (56/86 [65%] with *CDKN2A* deletion; 3/414 [1%] without *CDKN2A* deletion; Fisher’s exact *p* < 0.001; **figure 6E**). *MTAP*-deleted tumors had significantly higher MetScores than tumors without detected *MTAP* deletion (median [IQR], 0.055 [0.040] versus 0.024 [0.054]; Wilcoxon rank-sum *p* < 0.001; **figure 6F**). This association appeared to be largely attributable to *CDKN2A* co-deletion (**figure 6G**). In a multivariable linear model with Z-standardized MetScore as the outcome and both deletion indicators as predictors, *CDKN2A* deletion remained associated with higher MetScore after adjustment for *MTAP* deletion (adjusted mean difference, 0.72 SD; 95% CI, 0.39–1.06; *p* < 0.001). In contrast, *MTAP* deletion was not associated with MetScore after adjustment for *CDKN2A* deletion (adjusted mean difference, 0.17 SD; 95% CI, −0.23 to 0.56; *p* = 0.40). The fraction of *MTAP*-deleted and *MTAP-* preserved samples classified as basal subtype was similar (14/59 (23.7%) of deleted samples; 81/441 (18.4%) of intact samples; Fisher’s exact *p* = 0.38; **figure 6H**).

Collectively, these results demonstrate that the metastasis-high cell state is enriched in patient populations vulnerable to emerging targeted therapies including KRAS and PRMT5 inhibitors.

## DISCUSSION

In this study, we used the KYT cohort and an independent institutional cohort to validate and extend our previous observations regarding the metastatic-potential axis quantified by MetScore. MetScore was higher in metastatic-site biopsies than in primary tumors across two commercial RNA-sequencing platforms. Within the KYT cohort, this association was observed across sites typically seeded through hematogenous or lymphatic dissemination and was most pronounced in liver metastases. In contrast, peritoneal metastases had lower MetScores than primary tumors. Because PDAC cells can reach the peritoneum through transcoelomic dissemination without completing all steps required for vascular spread, these findings support distinct biological requirements for systemic and peritoneal colonization. They do not establish the underlying mechanisms, which will require experimental models of route-specific dissemination.

MetScore and basal subtype were each associated with shorter overall survival when modeled together, establishing that they provide prognostic information independent of one another. These associations were consistent across primary and metastatic-site biopsies and among patients with M0 and M1 disease at diagnosis, with no evidence of effect modification by biopsy site or metastatic status. The association between MetScore and survival in patients with M1 disease is particularly notable because it indicates that the biological programs captured by MetScore remain clinically relevant after metastatic disease has been established. Together with the dissemination-site findings, these results provide strong validation of metastatic potential as a clinically relevant dimension of PDAC transcriptomic state that is distinct from classical–basal differentiation.

Exploratory analyses further identified MetScore as a candidate modifier of the association between chemotherapy backbone and objective response. Among patients with MetScore-high tumors, 5-FU-based therapy was associated with a higher ORR than gemcitabine-based therapy, whereas no corresponding difference was detected among patients with MetScore-low tumors; this heterogeneity was supported by a treatment-by-MetScore interaction. Classical tumors also had a numerically higher ORR with 5-FU–based therapy, consistent with prior reports, and classical/MetScore-high tumors exhibited the largest observed treatment-associated difference in ORR. Thus, these data nominate classical/MetScore-high PDAC as a 5FU-response-enriched state. Given the nonrandomized design, heterogeneity of the treatment groups, and use of ORR as an endpoint, these findings require prospective validation in a randomized comparison of standardized regimens powered for a treatment–biomarker interaction using progression-free or overall survival. Because approximately 50% of patients with advanced PDAC do not remain eligible for second-line treatment^26^, selection of an effective first-line chemotherapy regimen is of critical importance. Similar transcriptome-based biomarker analyses could eventually be incorporated into studies combining or sequencing KRAS-directed therapies with chemotherapy and could be integrated with emerging artificial intelligence-guided histopathology-based biomarkers^27^.

The associations between transcriptomic state and TME architecture provide a potential biological context for the response findings. Higher MetScore was associated with a relative shift from stromal toward immune expression signals, whereas basal-state probability showed the opposite association. At finer resolution, MetScore was negatively associated with the MCP-counter fibroblast score, while basal-state probability was positively associated with this score; accordingly, classical/MetScore-high tumors had the lowest fibroblast-associated signal. Deep-learning-based analysis of matched H&E images showed a concordant reduction in connective/soft-tissue cells, providing orthogonal support for reduced nonimmune stromal cellularity. The co-occurrence of reduced fibroblast-associated signals and greater response enrichment with 5-FU-based therapy raises the hypothesis that stromal architecture contributes to the clinical association. Consistent with the need to consider non-cell-autonomous mechanisms, MetScore was not associated with sensitivity to either 5-FU or gemcitabine in isolated murine PDAC cells *in vitro*. Studies in colorectal cancer have shown that CAFs can promote 5-FU resistance through DKK1-mediated immunosuppression^28^ and COL8A1–ITGB1 interaction-induced epithelial-to-mesenchymal transition^29^, but whether these mechanisms operate in PDAC remains unknown. MetScore and basal-state probability also showed concordant associations with monocytic-lineage abundance. However, the use of bulk RNA sequencing and computational cell-population estimates limited cellular resolution. Single-cell and spatial studies in anatomically defined cohorts will be required to resolve the specific CAF and myeloid sub-populations associated with each transcriptomic axis.

In summary, this study establishes the metastatic-potential axis as a clinically relevant dimension of PDAC transcriptomic state that is distinct from classical–basal differentiation. The strongest findings demonstrate the generalizability of MetScore across real-world cohorts and sequencing platforms, its association with systemic rather than transcoelomic dissemination, and its prognostic relevance across disease settings. Exploratory analyses additionally nominate classical/MetScore-high PDAC as a 5-FU response-enriched state and identify stromal features that may contribute to this association. Together, these results provide a two-axis framework for prospective clinical validation and mechanistic studies aimed at intercepting systemic metastatic colonization and its interactions with the tumor microenvironment.

## ACKNOWLEDGEMENTS

We gratefully acknowledge the patients who participated in the Pancreatic Cancer Action Network Know Your Tumor program, whose participation and data made this research possible. We thank Drs. Shayan Monabbati and Satish Viswanath for their feedback on machine learning–based cell-type annotation of histopathology images. We also thank Greg Joseph of the Winship Data and Technology Applications (WinDATA) Shared Resource for identifying eligible patients for the Emory validation cohort and extracting the corresponding RNA-sequencing data.

## SUPPLEMENTAL INFORMATION

### Supplemental Document 1. Supplemental materials and methods

**Table S1. Chemotherapy regimens**.

**Table S2. Cell line MetScores**.

## TABLE AND FIGURE TITLES AND LEGENDS

**Figure S1.**
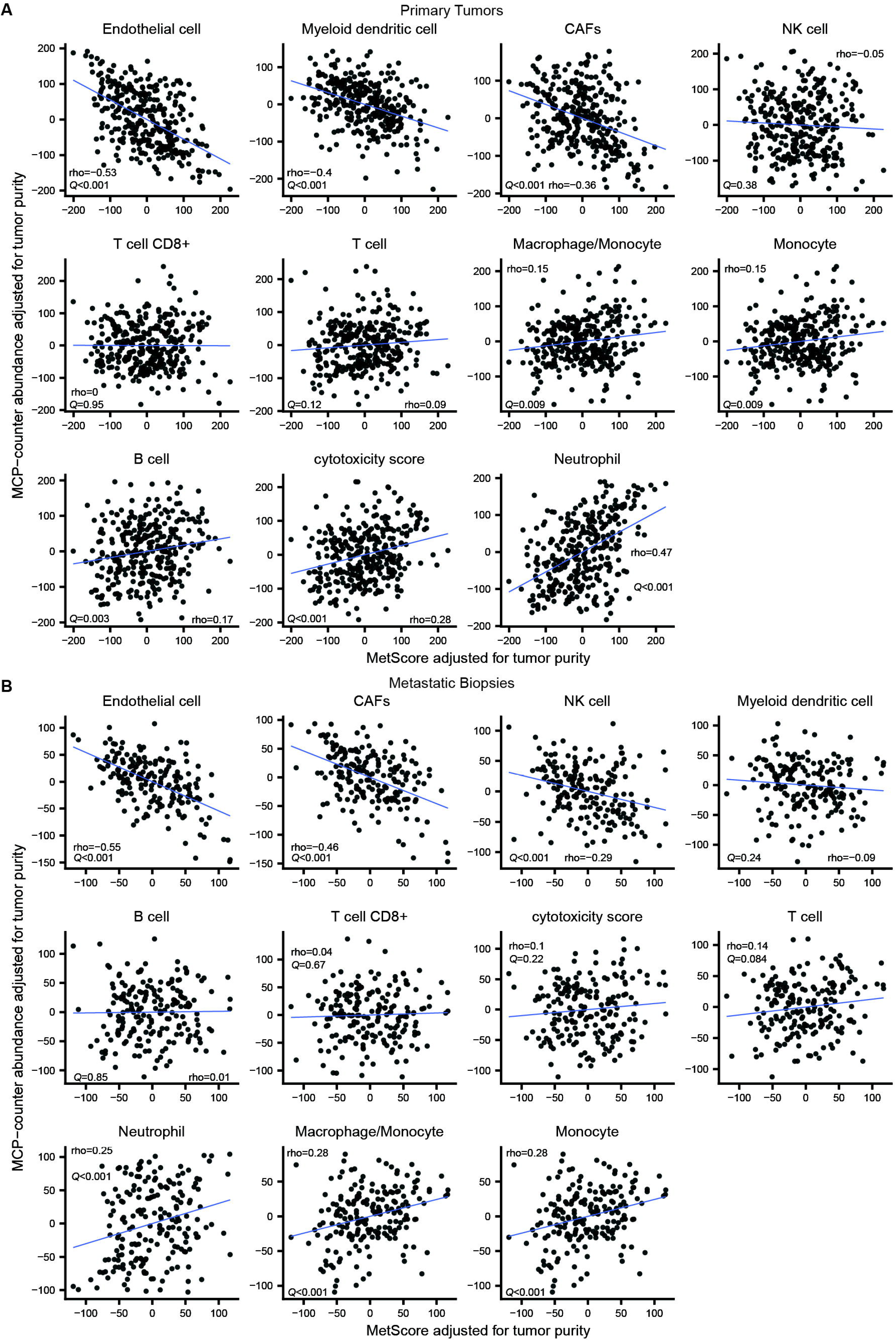
Related to Figure 5: MetScore-high and basal states have opposing associations with fibroblast abundance but concordant associations with monocytic-lineage cells. Associations between MetScore and MCP-counter cell-population scores tumors in the Know Your Tumor (KYT) cohort with available transcriptomic data stratified by anatomic site (primary tumors in **A**, n = 318; metastatic biopsies in **B**, n = 194). MetScore and MCP-counter scores were residualized for ESTIMATE-derived tumor purity before analysis. Each point represents one tumor biopsy. Least-squares regression lines are displayed for visualization, and Spearman’s ρ and corresponding *P* values are shown.

**Figure S2.**
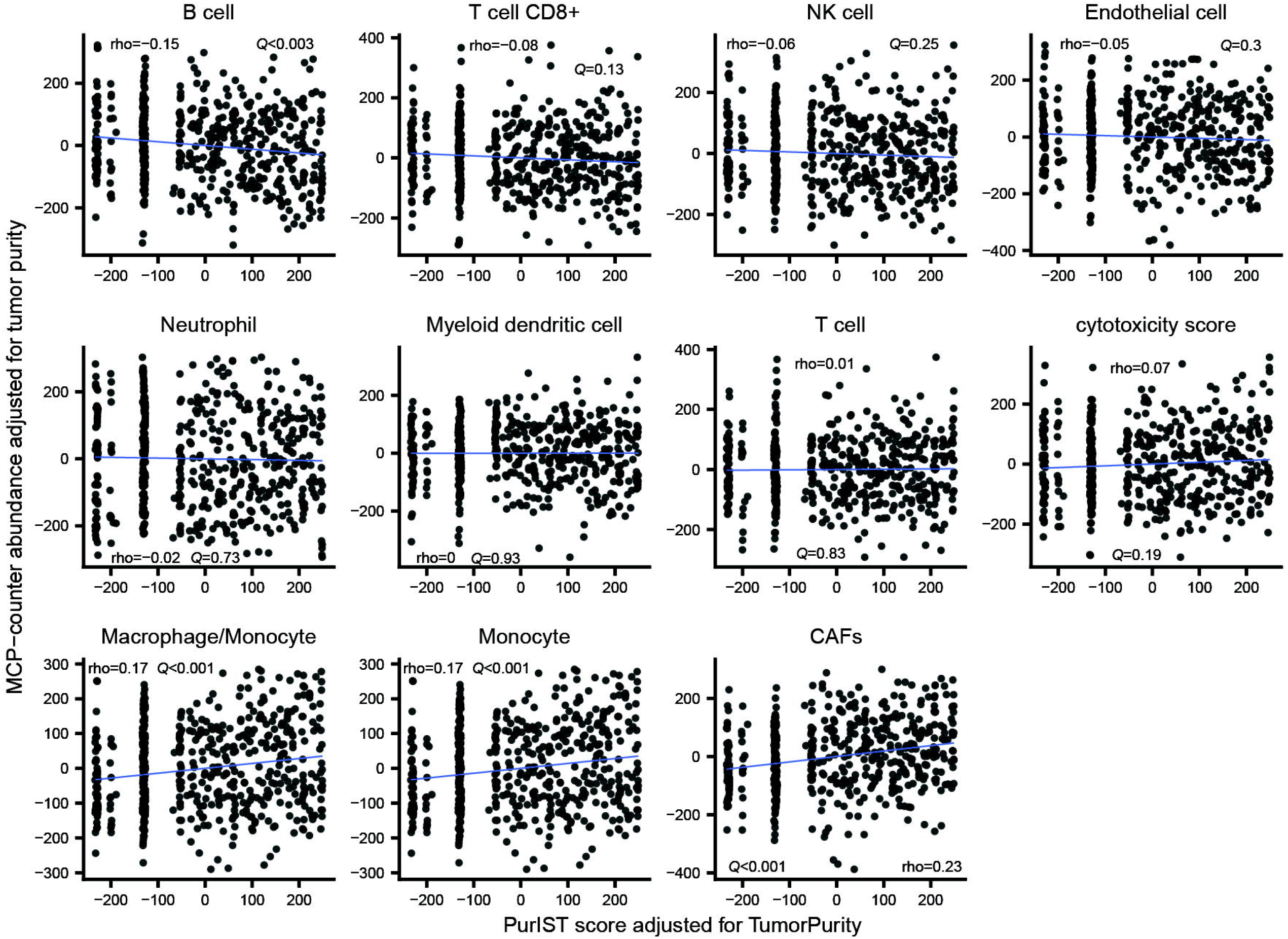
Related to Figure 5: MetScore-high and basal states have opposing associations with fibroblast abundance but concordant associations with monocytic-lineage cells. Associations between PurIST basal subtype probability score and MCP-counter cell-population scores tumors in the Know Your Tumor (KYT) cohort with available transcriptomic data (n = 512). PurIST and MCP-counter scores were residualized for ESTIMATE-derived tumor purity before analysis. Each point represents one tumor biopsy. Least-squares regression lines are displayed for visualization, and Spearman’s ρ and corresponding *P* values are shown.

**Figure S3.**
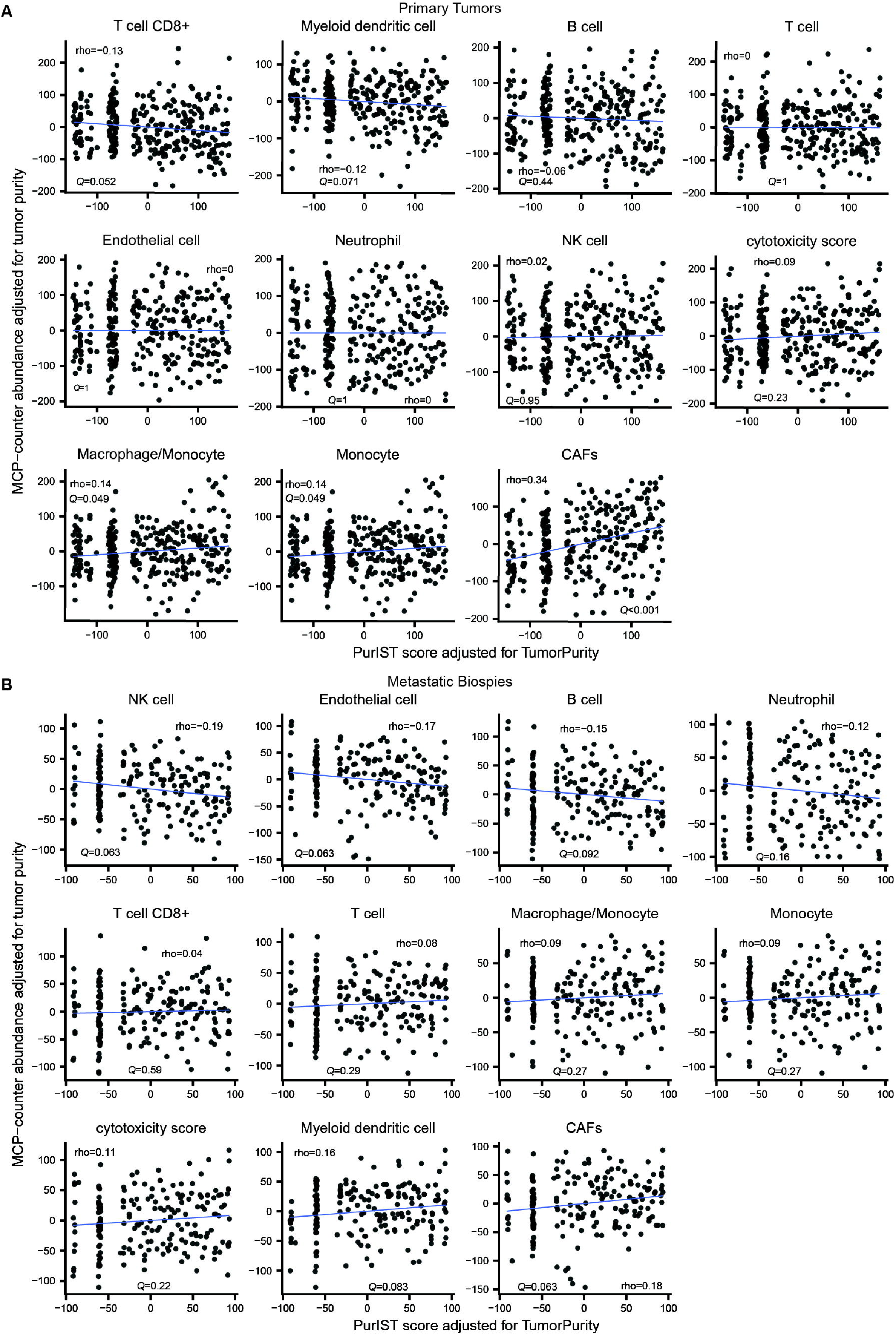
Related to Figure 5: MetScore-high and basal states have opposing associations with fibroblast abundance but concordant associations with monocytic-lineage cells. Associations between PurIST basal subtype probability score and MCP-counter cell-population scores tumors in the Know Your Tumor (KYT) cohort with available transcriptomic data stratified by anatomic site (primary tumors in **A**, n = 318; metastatic biopsies in **B**, n = 194). PurIST and MCP-counter scores were residualized for ESTIMATE-derived tumor purity before analysis. Each point represents one tumor biopsy. Least-squares regression lines are displayed for visualization, and Spearman’s ρ and corresponding *P* values are shown.

