## Supplemental Document 1 for "MetScore and classical–basal subtype intersect to define chemotherapy response and tumor microenvironment architecture in pancreatic ductal adenocarcinoma"

### SUPPLEMENTAL MATERIALS AND METHODS

#### Know Your Tumor cohort

De-identified data used in this study derives from the Pancreatic Cancer Action Network's (PanCAN) SPARK platform[1]. At the time of data access (9 March 2026), the platform contained 1,149 biospecimens from 825 patients enrolled in the Know Your Tumor (KYT) molecular profiling program. We excluded 315 normal-tissue biospecimens and one unspecified biospecimen, leaving 833 tumor biospecimens. We subsequently excluded 36 biospecimens from tumors with a recorded primary site other than the pancreas, leaving 797 pancreatic tumor biospecimens, of which 587 had available RNA-sequencing data.

Additional eligibility criteria were applied to define the final PDAC cohort. We excluded samples without a clearly annotated anatomic biopsy site (n = 4); samples whose annotated biopsy sites introduced ambiguity regarding the primary diagnosis, including extrahepatic bile duct (n = 9), ampulla of Vater (n = 4), colon not otherwise specified (n = 1), appendix (n = 2), and stomach (n = 2); and samples with a histologic diagnosis other than adenocarcinoma, infiltrating duct carcinoma, or mucinous adenocarcinoma (n = 53). After application of these criteria, the final cohort comprised 512 tumor biopsies from 510 patients. There were two patients with multiple biopsies represented in the dataset (both had two biopsies); they were excluded from downstream patient-level analyses.

KYT data were accessed via a data use agreement between Emory University and the Pancreatic Cancer Action Network.

#### Winship cohort

Pancreatic cancer specimens submitted from the Winship Cancer Institute for clinical molecular profiling by Caris Life Sciences between 2024 and 2025 were retrospectively identified. The cohort comprised 145 specimens, including 87 primary tumors and 58 metastatic lesions. Gene-level expression values quantified as transcripts per million (TPM) were provided under a data-use agreement between Emory University and Caris Life Sciences.

#### Transcriptomic signature scoring and tumor-microenvironment inference

Gene-level expression data were generated using the Tempus xR RNA-sequencing assay for the KYT cohort, the Caris RNA-sequencing platform for the Winship cohort, and a previously described in-house RNA-sequencing pipeline for murine PDAC cell lines[2]. Expression was quantified as transcripts per million (TPM). Untransformed TPM values were used for MetScore, PurIST, ESTIMATE, and MCP-counter analyses. Within each expression cohort, genes with a mean expression below 1 TPM were excluded.

MetScore was calculated using the fixed gene sets and scoring procedure described previously[2]. Genes were then ranked within each sample according to their relative expression using rankGenes() from the R package singscore[3,4] (version 1.30.0). MetScore was calculated using simpleScore(), with the previously defined species-specific metastasis-high gene set (n = 202 genes for human; n = 207 genes for mouse) as the upregulated gene set and the metastasis-low gene set (n = 174 genes for human; n = 182 genes for mouse) as the downregulated gene set. Signature genes absent from an expression matrix because they were not detected or did not meet the expression threshold were omitted from scoring.

Basal-state probabilities and classical–basal subtype assignments were generated from TPM expression values using the publicly available PurIST classifier (Rashid et al., 2020). Tumors with a predicted basal probability greater than 0.5 were classified as basal; tumors with a probability less than or equal to 0.5 were classified as classical. The continuous predicted probability of basal classification is referred to as basal-state probability.

ESTIMATE[5] ImmuneScore, StromalScore, ESTIMATEScore, and tumor-purity estimates were calculated from untransformed TPM values using the R package estimate (version 1.0.13), with estimateScore() and platform = "illumina". To quantify immune–stromal balance, ImmuneScore and StromalScore were independently Z-standardized across the KYT cohort. The immune–stroma balance score was calculated as

$$Z_{ImmuneScore} - Z_{StromalScore}$$

such that higher values indicated greater immune relative to stromal representation.

MCP-counter[6] cell-population scores were calculated from bulk RNA-sequencing profiles using the mcp_counter method implemented in the R package immunedeconv (version 2.1.0). The input was a gene-by-sample matrix of untransformed TPM values annotated with HGNC gene symbols. MCP-counter outputs were interpreted as relative cell-population scores and not as absolute cell fractions.

Associations between MetScore and each MCP-counter cell-population score were evaluated using partial Spearman rank correlations adjusted for ESTIMATE-derived tumor purity. Partial correlation coefficients and corresponding $p$ values were calculated using pcor.test() from the R package ppcor[7] (version 1.1), with method = "spearman". Each cell population was evaluated separately. The reported $p$ values are nominal and not corrected for multiple hypothesis testing.

For visualization, MetScore, the corresponding MCP-counter score, and tumor purity were rank-transformed, with average ranks assigned to tied observations. Ranked MetScore and the ranked MCP-counter score were separately regressed against ranked tumor purity using ordinary least-squares regression. The resulting residuals were plotted against one another, and ordinary least-squares regression lines without confidence bands were added for visualization. Cell populations were ordered by their partial Spearman correlation coefficients. The same procedures were repeated with basal-state probability in place of MetScore.

**Survival analysis**

Overall survival was defined as the number of days from the earliest documented initial diagnosis date to death or last known follow-up. When multiple diagnosis records were available, the earliest date recorded in complete YYYY-MM-DD format was used. Patients with a recorded date of death were coded as having experienced an event, whereas patients without a recorded death were censored at their last known follow-up date. Survival intervals with an end date preceding the initial diagnosis date were treated as missing.

The association between MetScore and overall survival was evaluated using a multivariable Cox proportional-hazards model implemented with coxph() in the R package survival[8] (version 3.8-3). MetScore was Z-standardized within the survival-analysis cohort and modeled continuously; hazard ratios therefore represent the change in the hazard of death per 1-standard-deviation increase in MetScore. The model was adjusted for PurIST subtype, with classical tumors serving as the reference group. The proportional-hazards assumption was evaluated using scaled Schoenfeld residuals with cox.zph().

For visualization, MetScore was dichotomized at the median, with values above the median classified as high and values at or below the median classified as low. Kaplan–Meier curves were generated using survfit() for four groups defined jointly by PurIST subtype and MetScore category: classical–low, classical–high, basal–low, and basal–high. Survival distributions were compared using a global log-rank test implemented with survdiff(). All tests were two-sided.

**Therapy response analysis**

Line-of-therapy records were linked to response assessments using patient and line-of-therapy identifiers and subsequently linked to available transcriptomic data by patient identifier. Analyses were restricted to first-line treatments (line_of_therapy_number = 1) with an evaluable MetScore and an available response assessment.

Treatment regimens were assigned by exact matching of recorded agent combinations to prespecified categories. Prespecified fluorouracil- or capecitabine-containing regimens were classified as 5-fluorouracil (5-FU)–based, and prespecified gemcitabine-containing regimens were classified as gemcitabine-based. Regimens containing both drug classes and regimens not assigned to either category were classified as "Other." Therapeutic-response analyses were restricted to patients receiving first-line 5-FU–based or gemcitabine-based therapy. The complete regimen classification is provided in **Table S1**.

Complete or partial response was classified as an objective response, whereas stable or progressive disease was classified as no objective response. When multiple assessments were recorded for the same patient and regimen, the best observed response was retained according to the following ordered categories: progressive disease, stable disease, partial response, and complete response. After this procedure, each patient contributed one observation. Patients without an assessment that could be mapped to one of these four categories were excluded.

Objective response rates (ORRs) were calculated as the number of patients with a complete or partial response divided by the total number of response-evaluable patients in each treatment and molecular subgroup. ORRs were evaluated between 5-FU–based and gemcitabine-based therapy within MetScore-low and MetScore-high tumors, within classical and basal PurIST subtypes, and within the four groups jointly defined by MetScore and PurIST subtype. MetScore-low and MetScore-high groups were defined using the cohort-level median as described above. Exact 95% binomial confidence intervals for ORRs were calculated using binom.test(). Within-group comparisons of objective response between treatment categories were performed using two-sided Fisher exact tests.

Potential treatment-effect modification by MetScore group or PurIST subtype was evaluated using binomial logistic-regression models. For each molecular classification, a model containing treatment and molecular-group main effects was compared with a model additionally containing their interaction using a likelihood-ratio test. Gemcitabine-based therapy, MetScore-low status, and classical subtype served as the reference categories.

To evaluate whether jointly considering MetScore and PurIST subtype provided additional treatment-predictive information, a model containing the treatment-by-MetScore interaction, subtype main effect, and MetScore-by-subtype interaction was compared with a full model containing the treatment-by-MetScore-by-subtype interaction and all corresponding lower-order terms. A separate likelihood-ratio test compared the full three-way-interaction model with a model containing all main effects and two-way interactions, thereby testing the three-way interaction specifically. All statistical tests were two-sided.

**In vitro drug-sensitivity analysis**

To assess whether MetScore was associated with cancer cell-intrinsic sensitivity to 5-fluorouracil (5-FU) or gemcitabine within the classical subtype, we used five previously established murine classical PDAC cell lines spanning a range of MetScores[2]. Cell-line identities and corresponding MetScores are provided in **Table S2**.

Cells were seeded into 96-well plates at 5,000 cells per well and allowed to adhere overnight (~18 hours). Cells were subsequently treated with 5-FU (MedChemExpress HY-90006) or gemcitabine (MedChemExpress HY-17026) across 8-point concentration ranges of 137nM-300μM and 4.6nM-10μM for 5-FU and gemcitabine, respectively, using 3-fold serial dilutions. Vehicle-treated (0.1% DMSO) and cell-free wells were included on each plate. After 72 hours of drug exposure, cellular viability was measured at 570nm using the CellTiter 96 Non-Radioactive Cell Proliferation Assay (Promega G4000) according to the manufacturer's instructions.

At each concentration, percent viability was calculated as:

$$\frac{\left(\overline{A}_{sample} - \overline{A}_{blank}\right)}{\left(\overline{A}_{vehicle} - \overline{A}_{blank}\right)} \cdot 100$$

where $\overline{A}$ denotes mean absorbance across three technical replicates. Normalized AUC was calculated by trapezoidal integration of the viability fraction over the log10-transformed concentration range and division by the width of that range. Thus, normalized AUC represents mean relative viability across the tested log-concentration range, with higher values indicating greater residual viability and lower drug sensitivity.

Three independent experiments were performed, each containing 3 technical replicates per cell line per concentration. Associations between MetScore and normalized AUC averaged across the three experiments were evaluated using two-sided Spearman correlations and exact permutation *p* values.

**CellViT-based cell segmentation and classification**

Digitized whole-slide images of hematoxylin and eosin-stained tissue sections from the KYT cohort were analyzed using the pretrained CellViT++-SAM-H model with using PanNuke nuclei classification (CellViT++ version 1.0.0b). Whole-slide images were processed using the authors' inference pipeline at 0.25 microns per pixel. Slides with less than 1,000 cells and greater than 10% dead cells were removed from downstream analysis.

CellViT was used to segment individual cells and classify them as neoplastic, non-neoplastic epithelial, inflammatory, connective tissue, or dead. For each tumor, the connective-tissue fraction was calculated as the number of cells classified as connective tissue divided by all classified cells. Tumors without an evaluable matched image were excluded from digital-pathology analyses, leaving 502 tumors.

##### Somatic genomic alterations

Somatic single-nucleotide variants, indels, and copy-number alterations were obtained from targeted DNA sequencing performed using the Tempus xT assay. SNV/indel analyses were restricted to tumors with at least one reportable SNV or indel call (n = 499), whereas copy-number analyses were restricted to tumors with *MTAP* and *CDKN2A* copy-number calls (n = 500).

Tumors with at least one reportable *KRAS* SNV or indel were classified as *KRAS*-mutant. Tumors with at least one reportable alteration elsewhere in the assayed panel but no detected *KRAS* alteration were classified as having no detected *KRAS* mutation. This designation indicates the absence of a reported *KRAS* alteration within the assay's reportable range and does not establish the absence of an unreported or subthreshold tumor alteration.

Three tumors contained two detected *KRAS* variants. In each case, one was a canonical hotspot mutation and the other was a non-hotspot variant. For the mutually exclusive allele categories used in Figures 6A&C, these tumors were classified according to the canonical hotspot mutation. In Figure 6B, all three tumors were placed in the *KRAS*-mutant category.

*MTAP* and *CDKN2A* deletion status was determined exclusively from reportable gene-level copy-number calls. A tumor was classified as deletion-positive when the gene-level record had an estimated copy number of 0.

The overlap between MTAP and CDKN2A deletion was evaluated using a two-sided Fisher exact test, with an odds ratio and 95% confidence interval reported. To assess their conditional associations with MetScore, Z-standardized MetScore was modeled as the outcome in a multivariable linear-regression model containing both MTAP and CDKN2A deletion indicators. Tumors were also descriptively classified into four mutually exclusive groups: neither deletion, CDKN2A deletion only, MTAP deletion only, and MTAP/CDKN2A co-deletion.

##### Patient and Public Involvement

This study was conducted in collaboration with the Pancreatic Cancer Action Network (PanCAN), a pancreatic cancer patient advocacy organisation that operates the Know Your Tumor programme. PanCAN staff members contributed as coauthors and provided feedback on the manuscript. PanCAN will support dissemination of the findings directly to patients. Planned dissemination also includes presentation of the work at the 2026 PanCAN symposium.
